# Energetically constrained turnover drives the emergence of aging

**DOI:** 10.64898/2026.05.19.726278

**Authors:** Misana Yao, Shinji Deguchi

**Affiliations:** Division of Bioengineering, Graduate School of Engineering Science, The University of Osaka; Global Center for Medical Engineering and Informatics, The University of Osaka; R^3^ Institute for Newly-Emerging Science Design, The University of Osaka

## Abstract

Aging is characterized by progressive functional decline, yet why such decline is observed broadly across living systems remains unclear. While molecular and cellular mechanisms describe how aging progresses, they do not explain why functional decline should arise as a natural consequence of living organization. Here, we show that aging naturally emerges from three general features of life: unavoidable damage, turnover-mediated maintenance, and the energetic constraint of turnover. We develop a hierarchical damage–turnover model in which component-level damage and energetically constrained turnover jointly determine whole-system performance. In the model, damage stochastically converts functional components into non-functional components, whereas turnover restores component performance at a rate coupled to whole-system performance. Analytical and Monte Carlo analyses reveal two regimes: a non-aging regime, in which performance remains finite, and an aging regime, in which performance progressively collapses toward zero. Performance-independent turnover always maintains a positive steady state, whereas performance-dependent turnover generates irreversible decline when reduced performance weakens maintenance capacity. Stochastic fluctuations further promote collapse near the transition boundary, even when deterministic analysis predicts a nonzero steady state. These results indicate that unavoidable damage and energetically constrained turnover are sufficient to generate aging-like decline, providing a minimal theoretical explanation for long-term irreversibility in biological systems.

## Introduction

Aging is characterized by a progressive decline in biological function over time (López-Otín et al., 2013). Although the specific features of aging vary across forms of life, this functional decline is broadly observed in animals, plants, fungi and even unicellular organisms. In animals, aging is often associated with reduced immune competence and physical performance (Nussey et al., 2013; Peters et al., 2019), whereas in plants it can involve changes in growth and regenerative capacity (Ye et al., 2020; Yun, 2015). Aging-like phenomena have also been reported in fungi and unicellular organisms as alterations in replicative capacity (Bhattacharya et al., 2021; Florea, 2017). Moreover, aging is not restricted to whole organisms but also occurs at the levels of individual cells, tissues, and organs (Di Micco et al., 2021; Tchkonia et al., 2010; Wyss-Coray, 2016). Thus, despite its diverse manifestations, aging is characterized by the progressive deterioration of biological function and can be regarded as a fundamental property of living systems.

As aging progresses, the gradual deterioration of biological function increases susceptibility to a wide range of age-related diseases, including cancer, cardiovascular disease, metabolic disorders, and neurodegenerative diseases (Guo et al., 2022; Niccoli & Partridge, 2012). Aging represents a major risk factor for these conditions, and many leading causes of death are associated with age-related pathological changes (Kennedy et al., 2014). Elucidating the fundamental biological mechanisms underlying aging is therefore essential for understanding how functional decline gives rise to disease vulnerability and for developing more effective strategies to prevent age-related diseases.

Extensive studies have identified numerous molecular and cellular processes associated with aging, including genomic instability, telomere attrition, epigenetic alterations, loss of proteostasis, impaired autophagy, deregulated nutrient sensing, mitochondrial dysfunction, cellular senescence, stem cell exhaustion, altered intercellular communication, chronic inflammation, and microbiome dysbiosis (Chantachotikul et al., 2025; Liu et al., 2022; López-Otin et al., 2023). While these studies have provided detailed descriptions of the processes through which aging progresses and have contributed substantially to therapeutic and diagnostic development, they do not by themselves explain why aging emerges across living systems or why it appears to be an inevitable consequence of biological organization. It also remains unclear whether aging is intrinsically irreversible or can, at least in principle, be fundamentally reversed.

Several studies have addressed the inevitability, irreversibility, and underlying mechanisms of aging. However, many of these studies have focused on specific aging-related phenomena (Karin et al., 2019; Nelson & Masel, 2017; Wodarz, 2007), leaving open the question of whether aging can be explained from more general principles. In this study, we theoretically examine how aging can arise as an inevitable process from minimal structural and dynamical features shared by living systems. Specifically, we consider a system composed of constituent components and the whole formed by those components, together with damage–repair dynamics at the component level. Using this framework, we investigate the conditions under which aging emerges as an inevitable process and the extent to which it can, in principle, be reversed. Our results show that aging can arise from the coupling between component-level damage–repair dynamics and whole-system function, while also identifying conditions under which this inevitability may be avoided.

## Methods

### Model overview

We construct a minimal two-layer model to examine how aging can emerge from the coupling between component-level damage–turnover dynamics and whole-system performance. The model consists of constituent components and the whole system formed by those components (Fig.1). This abstraction represents the simplest hierarchical relationship in which the performance of the whole depends on the performance states of its components.

**Fig.1.**
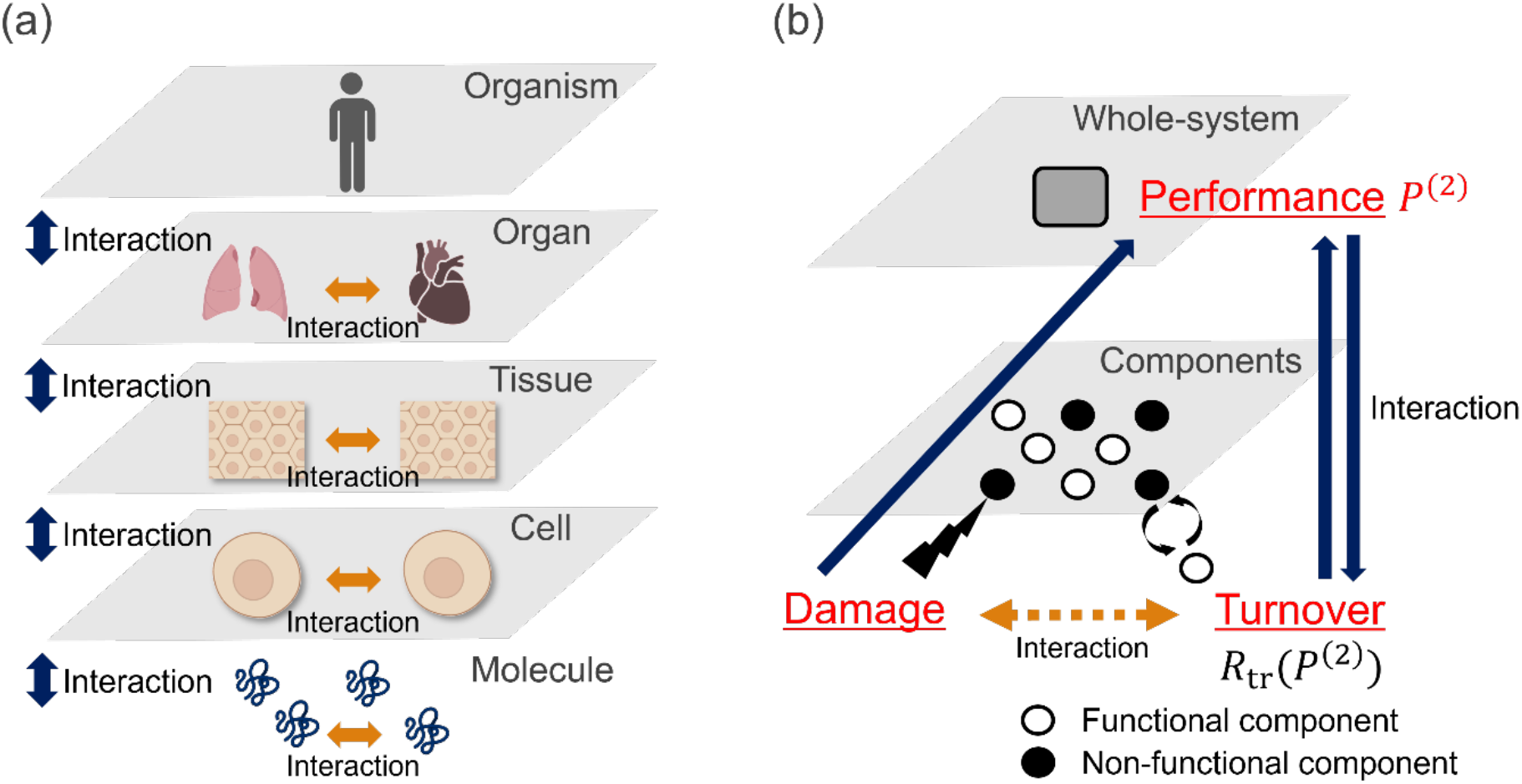
Hierarchical structure and interactions. **(a)** Examples of hierarchical organization in biological systems. Interactions occur both among elements within a layer and across different hierarchical levels. **(b)** The two-layer abstraction used in the model, consisting of constituent components and the whole-system formed by these components. Direct interactions are assumed between layers, whereas interactions among components arise indirectly through the influence of the higher layer.

The model is based on three broadly observed features of living systems. First, components are continuously exposed to intrinsic and extrinsic damage, including DNA replication errors, oxidative byproducts of metabolism, ultraviolet radiation, and other environmental insults (López-Otín et al., 2013; Panich et al., 2016). Second, living systems maintain function through turnover processes that repair damaged components or replace existing damaged components with new functional ones (Reddien, 2024; Saito & Deguchi, 2023; Toyama & Hetzer, 2013). These processes include molecular-level molecular-level maintenance of proteostasis through protein turnover and autophagy (Hernández-Cáceres et al., 2021; Mizushima & Press, 2011; Matsui et al., 2018; Saito et al., 2022), as well as cellular turnover in proliferative tissues (Pellettieri & Alvarado, 2007; Saw et al., 2017). Third, turnover is not merely a passive process but an actively regulated and energetically costly maintenance function. This property is exemplified by autophagy, which is executed through coordinated signaling pathways and ATP‐consuming molecular machinery rather than arising spontaneously. Indeed, the loss of essential autophagy-related factors results in embryonic lethality, demonstrating that turnover-like processes are tightly controlled and indispensable for maintaining cellular homeostasis (Mizushima & Levine, 2010; Tsukamoto et al., 2008). Therefore, turnover capacity is linked to whole-system performance (Fig.1).

In the model, each component is represented by a binary performance state, and whole-system performance is defined as the average performance of all components. Damage converts functional components into non-functional components, whereas turnover restores component performance. The coupling between component-level damage–turnover dynamics and performance-dependent maintenance capacity at the whole-system level constitutes the core structure of the model.

### Definition of component and whole-system performance

First, we model the hierarchical organization of biological systems using this minimal two-layer structure. The lower layer consists of *C* components, where *C* is treated as a constant, and the upper layer represents the whole system formed by these components. To quantify functional deterioration associated with aging, we introduce performance variables for both the components and the whole system. Each component *i* is assigned a binary component-level performance value 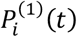, which indicates whether the component is functional at time *t*:

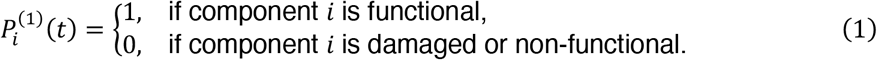

The whole-system performance *P*^(2)^(*t*) is then defined as the average of the component-level performance values:

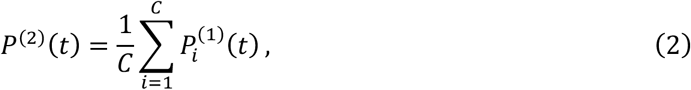

where *C* denotes the total number of components in the system. Thus, *P*^(2)^(*t*) takes continuous values between 0 and 1, with lower values corresponding to greater functional deterioration of the whole system.

### Damage and turnover dynamics

Damage and turnover are modeled as stochastic processes occurring at the component level (Fig.2). Damage occurs with rate *R*_d_ per unit time for each component and converts a functional component into a non-functional component. Turnover occurs with rate *R*_tr_ per unit time for each component and restores the component to the functional state. Although the occurrence of turnover events is treated probabilistically in time, turnover itself represents an actively regulated maintenance process that requires biological activity and energetic investment. Given these rates, the expected numbers of damaged and turned-over components per unit time are *CR*_d_ and *CR*_tr_, respectively, for *C* components. For generality, damage and turnover events are applied independently of the prior state of each component. Thus, a component can be selected for damage even if it is already non-functional, and a component can undergo turnover even if it is already functional.

**Fig.2.**
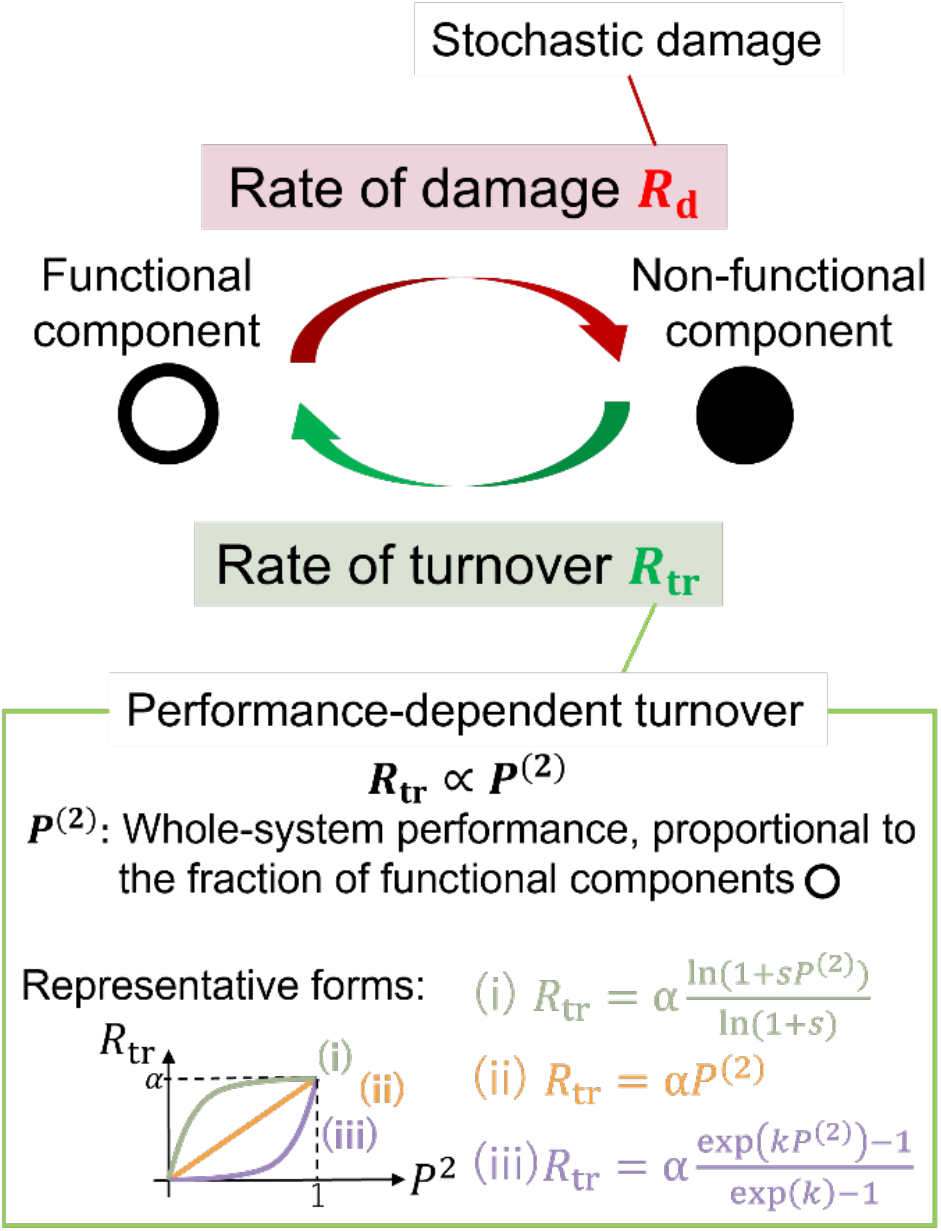
Overview of the dynamics of damage and turnover. Damage accumulates stochastically in each component, whereas turnover occurs with a rate that depends on the whole-system performance. Three representative forms of performance-dependent turnover are considered: **(i)** a logarithmic response, **(ii)** a linear response, and **(iii)** an exponential response. The parameter *α*denotes the maximum turnover rate; the parameter *s* controls the curvature of the logarithmic function; and the parameter *k* controls the steepness of the exponential function.

### Performance-dependent turnover

We next specify the dependence of the turnover rate on whole-system performance. Because turnover represents an actively regulated and energetically costly maintenance process, the capacity for turnover is taken to increase with whole-system performance *P*^(2)^(*t*). We denote this performance-dependent turnover rate as *R*_tr_(*P*^(2)^(*t*)).

To examine how the shape of this dependence affects the system dynamics, we consider three representative functional forms (Fig.2): (i) a logarithmic form, in which the turnover rate increases rapidly at low *P*^(2)^ and then saturates; (ii) a linear form; and (iii) an exponential form, in which the turnover rate increases slowly at low *P*^(2)^ and accelerates at high *P*^(2)^. In all cases, the turnover rate satisfies the boundary conditions *R*_tr_(0) = 0 and *R*_tr_(1) = *α*, where *α* ∈ [0,1] represents the maximum turnover rate. These boundary conditions represent that turnover cannot be sustained when whole-system performance is completely lost, whereas turnover reaches its maximum capacity when the whole system is fully functional.

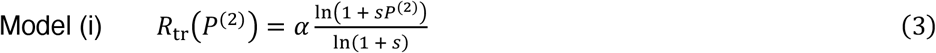

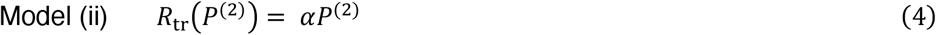

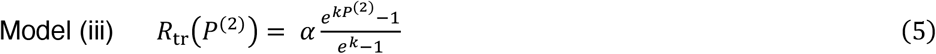

Here, *s* controls the curvature of the logarithmic response, with larger *s* making the turnover rate more sensitive to changes in *P*^(2)^ at low performance. The parameter *k* controls the steepness of the exponential response, with larger *k* producing a more pronounced increase in turnover rate at high *P*^(2)^.

The resulting model dynamics are determined by stochastic damage and turnover events at the component level and by the dependence of turnover rate on whole-system performance. Initially, all *C* components are in the functional state, such that 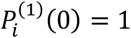 for all *i*, and therefore *P*^(2)^ = 1. At each time step, components are selected for damage with rate *R*_d_, which converts them to the non-functional state. Components are also selected for turnover with rate *R*_tr_(*P*^(2)^7, which restores them to the functional state. Through these stochastic transitions, the dynamic states of individual components collectively determine whole-system performance, which in turn modulates the turnover rate of the components. In this way, the model represents a structure in which direct interactions arise between the components and the whole system, while indirect interactions emerge among components through their shared dependence on whole-system performance.

Here, aging is defined as the progressive decline of whole-system performance *P*^(2)^(*t*) over time, resulting from the dynamics of damage and turnover. When all components lose their performance and *P*^(2)^(*t*) reaches zero, turnover can no longer be executed because *R*_tr_(0) = 0. Under this condition, the system loses its capacity for recovery and is regarded as dead.

### Governing equation and analytical solution

To analytically characterize the behavior of the model, we first derive the governing equation for whole-system performance. For analytical convenience, the whole-system performance *P*^(2)^(*t*) is denoted as *P*(*t*) from this section. Within an infinitesimal time interval Δ*t*, damage events decrease *P*(*t*) in proportion to the fraction of components that remain functional, which is given by *P*(*t*) itself. In contrast, turnover events increase *P*(*t*) in proportion to the fraction of non-functional components, 1 − *P*(*t*) by restoring non-functional components to the functional state. Because damage and turnover can occur within the same time interval Δ*t*, we assume that damage occurs first and turnover follows. Based on this assumption, the temporal update of whole-system performance is written as

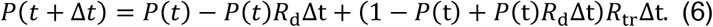

Taking Δ*t* sufficiently small allows us to neglect second-order infinitesimal terms, yielding

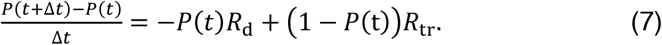

In the limit Δ*t* → 0, we obtain the governing equation

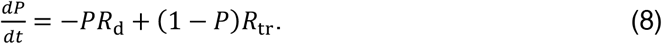

Among the three models, only the linear-form Model (ii) allows an analytical solution. Substituting this relation into the governing equation yields

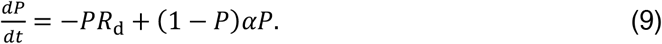

Solving this equation with the initial condition *P*(0) = 1, we obtain

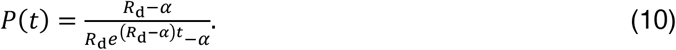

The equilibrium behavior of the linear turnover model is characterized by solving the fixed-point condition *dP*/*dt* = 0. The equilibrium value of whole-system performance is denoted as *P*^*^, and only solutions within the admissible range 0 ≤ *P*^*^ ≤ 1 are considered. This analysis shows that the system converges to *P*^*^ = −(*R*_d_ − *α*)/*α* when *α* > *R*_d_, whereas it converges to *P*^*^ = 0 when *α* < *R*_d_ (Fig.3(a)). Thus, the long-term state of the system is determined by the balance between the damage rate *R*_d_ and the maximum turnover rate *α*.

**Fig.3.**
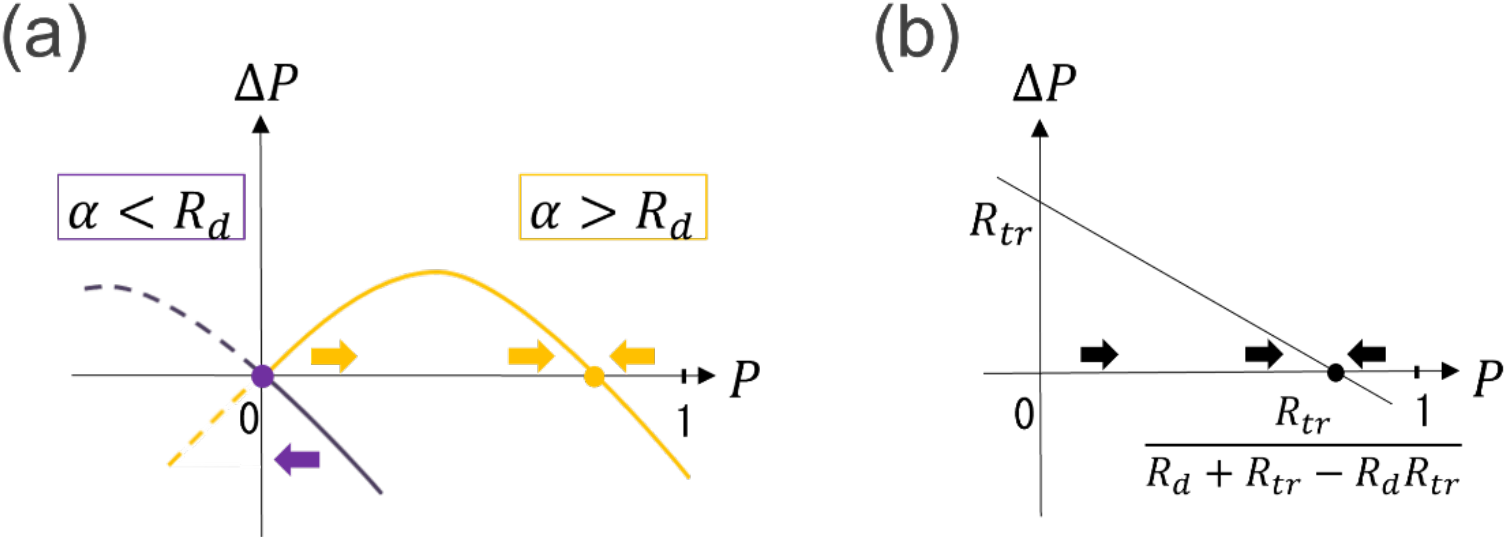
Numerical calculation of stability analysis. **(a)** The result of performance-dependent model where turnover rate is given *R*_tr_ = *αP*. The blue line indicates the case where *α* > *R*_d_, and the orange line indicates the case where *α* < *R*_d_. **(b)** The result of performance-independent model where turnover rate is given *R*_tr_ = *P*.

### Performance-independent turnover model

To examine the specific contribution of performance-dependent turnover, which constitutes a key mechanism of the present model, we analyze a simplified reference model in which turnover does not depend on whole-system performance. Here, the turnover rate is treated as a constant and set to *R*_tr_ = *α*. The time evolution of whole-system performance is then described by

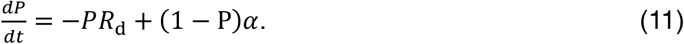

Solving this equation with the initial condition *P*(0) = 1 yields

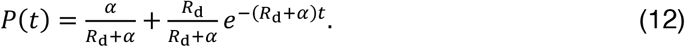

The fixed-point condition *dP*/*dt* = 0 gives a unique equilibrium value *P*^*^ = *α*/(*R*_*d*_ + *α*) for any *R*_d_ and *α*. Thus, in contrast to the performance-dependent turnover model, whole-system performance always converges to a positive equilibrium value in this case (Fig.3(b)).

### Numerical simulations

In addition to the analytical treatment, we performed Monte Carlo simulations for Models (i)–(iii) and for the performance-independent turnover model. Numerical simulations were used to analyze models that do not admit analytical solutions and to investigate the effects of stochastic fluctuations and parameter dependence on the system dynamics.

At the beginning of each simulation, all *C* components were initialized in the functional state 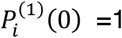 for all *i*, with *C* = 100). Time was discretized with a step size Δ*t* = 0.2, which was selected to approximate the continuous-time dynamics and to reproduce the analytical solutions in solvable cases. In each time step, the probabilities of damage and turnover events for each component were given by *R*_d_Δ*t* and *R*_tr_(*P*^(2)^)Δ*t*, respectively. For each component, uniform random numbers generated using NumPy in Python were used to determine whether damage and turnover events occurred. Damage events were evaluated first, followed by turnover events, in accordance with the update rules defined above.

After all components were updated within a time step, whole-system performance *P*^(2)^(*t*) was computed according to Eq. (2). The turnover rate for the next time step was then updated based on the resulting whole-system performance. Each simulation was run for *T* time steps, typically *T* = 400/Δ*t* (Δ*t* = 0.2). For each parameter set, simulations were repeated *N* times, typically *N* = 100, to obtain averaged trajectories and reduce stochastic variability. In parameter-scan analyses, *R*_d_ and *α* were varied from 0.1 to 0.9 in increments of 0.1. Large-scale scans were performed with *N* = 50 to reduce computational cost. A system was classified as having reached a steady state if *P*^(2)^(*t*) did not vary by more than 0.02 over the interval from *t* = 200/Δ*t* to *t* = *T*/Δ*t*.

## Results

### Performance-dependent turnover generates distinct aging and non-aging regimes

To investigate how different forms of performance-dependent turnover influence aging, defined here as the decline of whole-system performance, we analyzed the temporal evolution of whole-system performance *P*(*t*) in four turnover models: (i) a logarithmic model in which the turnover rate increases rapidly at low *P* and then saturates, (ii) a linear model, (iii) an exponential model in which the turnover rate accelerates at high *P*, and a performance-independent turnover model used as a reference. For each model, *P*(*t*) was evaluated using Monte Carlo simulations under identical parameter sets. In addition, for the linear and performance-independent turnover models, analytical solutions for *P*(*t*) were derived from the governing equations and compared with the numerical simulations.

Across all models, two qualitatively distinct dynamical regimes were observed: (A) a non-aging regime, in which *P*(*t*) converges to a non-zero equilibrium value in the analytical solution or fluctuates around that value in the numerical simulations, and (B) an aging regime, in which *P*(*t*) progressively declines toward zero (Fig.4). The logarithmic model (Model (i)) predominantly exhibited regime (A), whereas the exponential model (Model (iii)) consistently exhibited regime (B). In contrast, the linear model (Model (ii)) displayed a clear transition between the two regimes depending on parameter values: regime (A) emerged when *α* > *R*_*d*_, whereas regime (B) appeared when *α* < *R*_*d*_. The performance-independent turnover model exhibited only regime (A) across all examined parameter sets.

**Fig.4.**
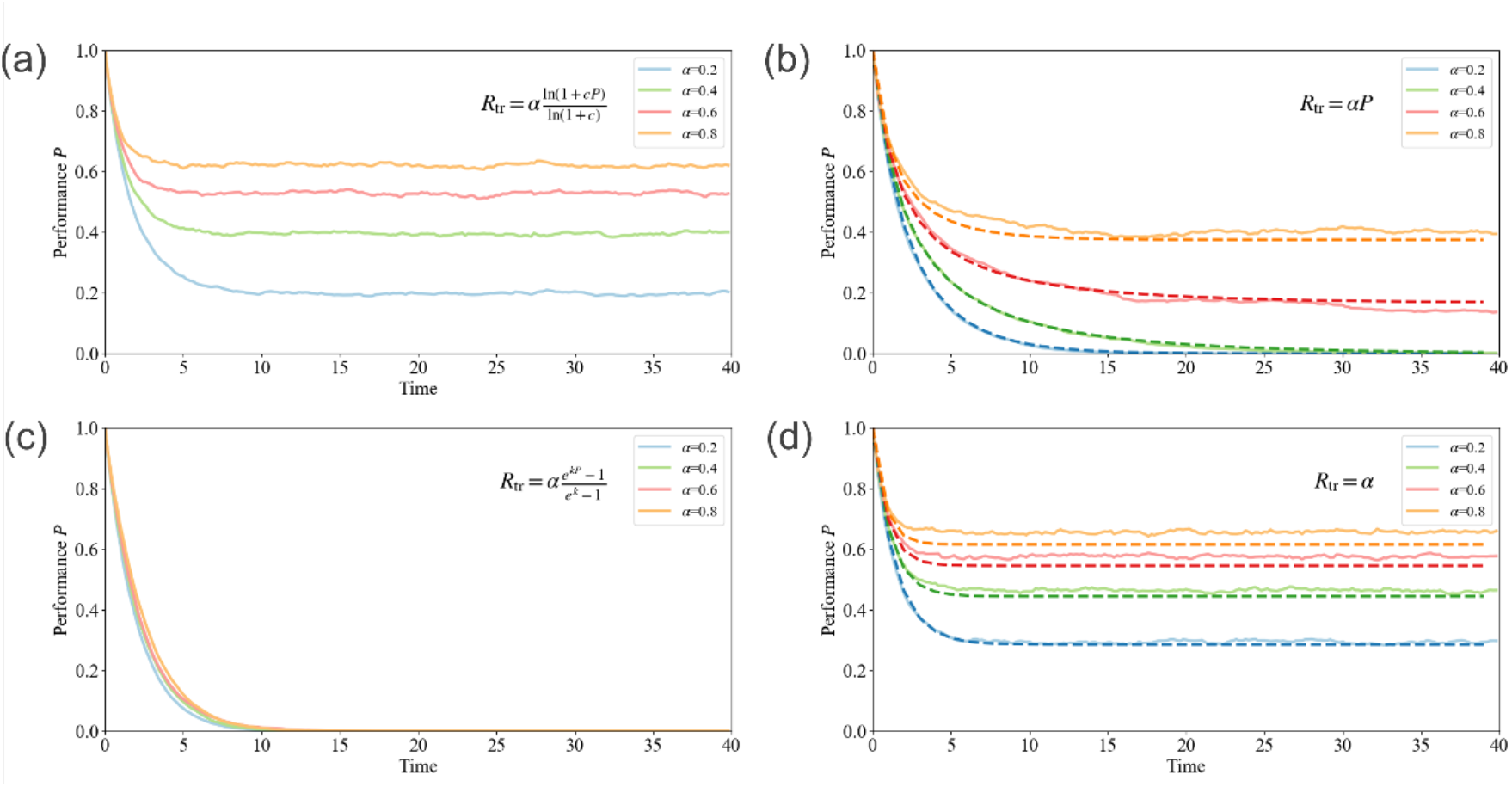
Temporal evolution of the whole-system performance *P*(*t*). **(a–d)** Numerical simulation results are shown as solid lines, and analytical solutions are shown as dashed lines in panels **(b)** and **(d). (a)** Results for the logarithmic turnover model (case (i)), with the curvature parameter fixed at *s* = 50. **(b)** Results for the linear turnover model (case (ii)), where the analytical solution follows the governing equation(10). **(c)** Results for the exponential turnover model (case (iii)), with the steepness parameter fixed at *k* = 4. **(d)** Results for the performance-independent turnover model, for which the analytical solution also follows directly from the governing equation(12). In all panels, the damage rate is fixed at *R*_d_ = 0.5, and the maximum turnover rate *α* is varied across *α* = 0.2, 0.4, 0.6, and 0.8.

For the linear and performance-independent turnover models, the numerical simulations agreed with the analytical solutions derived from the governing equations (Fig.4(b), Fig.4(d)). In particular, the transition boundary observed in the linear model was consistent with the fixed-point analysis, indicating that the balance between the damage rate *R*_*d*_ and the maximum turnover rate *α* determines whether the system maintains a finite equilibrium performance or undergoes progressive collapse.

### Stochastic fluctuations generate variability in individual aging trajectories

To investigate how stochastic fluctuations influence the aging dynamics observed at the ensemble level, we next analyzed individual simulation trajectories for the linear turnover model under the same parameter conditions as in Fig.4 (Fig.5). While the ensemble-averaged trajectories exhibited smooth temporal changes in *P*(*t*), individual realizations showed repeated local increases and decreases in performance over time. These individual trajectories correspond to single stochastic realizations of the model and can be interpreted as representing variability across individual systems under identical parameter conditions.

**Fig.5.**
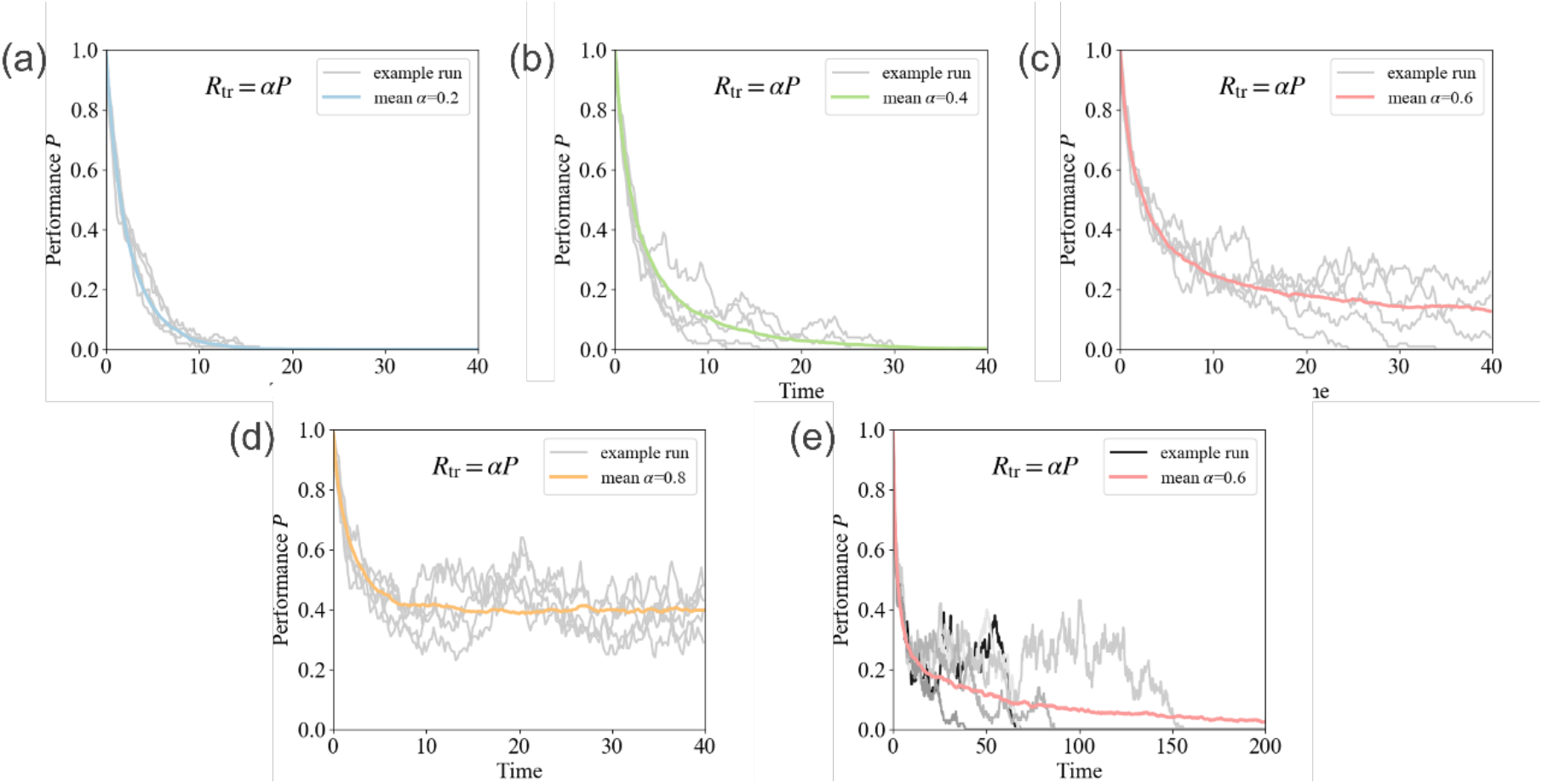
Sample individual trajectories of numerical simulation results for Model (ii) with *R*_*d*_ = 0.5. Panels **(a)**–**(d)** show representative simulation results for *α* = 0.2, 0.4, 0.6, and 0.8, respectively. Panel **(e)** displays the case *α* = 0.6over a longer time span for investigation. For each value of *α*, the colored line denotes the ensemble-averaged performance, while the gray lines represent individual trajectories. As an exception, in panel **(e)**, example runs are shown using grayscale lines for visual clarity.

These fluctuations arise from stochastic variability in the actual numbers of damaged and renewed components at each time step. Specifically, the net change in *P*(*t*) is determined by the difference between the number of components that become newly damaged and the number of components whose performance is restored through turnover. Because the component selection of both damage and turnover events occur probabilistically, this difference fluctuates over time, leading to transient positive or negative displacements in whole-system performance within individual trajectories.

For *α* = 0.2 and *α* = 0.4, these stochastic fluctuations produced substantial variability in the timing of performance collapse across trajectories. Although local increases in *P*(*t*) were intermittently observed, all trajectories exhibited an overall decline toward zero, consistent with the aging regime identified in the ensemble-average dynamics. In contrast, for *α* = 0.8, individual trajectories fluctuated around finite performance values without showing persistent drift toward collapse. Consequently, the ensemble-averaged dynamics stabilized around a nonzero steady-state value, corresponding to the non-aging regime.

A qualitatively distinct behavior emerged near the transition boundary at *α* = 0.6. Analytical calculations predict that this parameter set belongs to the non-aging regime because the equilibrium value remains positive. However, the ensemble-averaged trajectory exhibited an approximately linear decline over time, resembling the aging regime. To investigate this discrepancy, we examined individual trajectories over longer timescales (Fig.5(e)). Although each trajectory initially fluctuated around a finite value, occasional large damage events caused abrupt decreases in *P*(*t*). Once performance approached zero, turnover became insufficient to restore the lost performance because the turnover rate itself decreased with declining *P*(*t*). As a result, trajectories underwent irreversible collapse toward the dead state.

Because such stochastic collapse events can occur at any time step, the number of collapsed trajectories gradually accumulates across simulations. This accumulation produces an apparent linear decline in the ensemble-averaged performance, thereby giving rise to behavior resembling the aging regime. This behavior arises when the equilibrium point predicted by the analytical fixed-point solution lies close to zero. For the linear model, this equilibrium is given by *P*^*^ = 1 − *R*_*d*_/*α*, indicating that *P*^*^ becomes small when *α* only slightly exceeds *R*_*d*_. This interpretation is consistent with the simulation result in which this phenomenon appeared at *α* = 0.6 and *R*_*d*_ = 0.5. These results show that stochastic fluctuations at the single-trajectory level can alter the apparent ensemble-level aging dynamics, particularly near the transition boundary.

### Phase diagrams identify parameter regimes for aging and non-aging dynamics

Building on the previous analyses, we next systematically investigated the parameter space (*R*_d_, *α*) to determine under which conditions the aging and non-aging regimes emerge. For this purpose, phase diagrams were constructed for all turnover models based on Monte Carlo simulations (Fig.6). For Model (i), in which the turnover rate increases rapidly at low *P* and subsequently saturates, the non-aging regime predominated across most of the parameter space. The aging regime appeared only when the maximum turnover rate *α* was small relative to the damage rate *R*_d_. In contrast, Model (iii), in which the turnover rate accelerates at high *P*, exhibited the aging across the entire parameter space. Model (ii), in which the turnover rate depends linearly on *P*, exhibited an intermediate behavior between Models (i) and (iii). In particular, the aging regime always appeared when *α* ≤ *R*_d_, consistent with the analytical condition derived from the fixed-point analysis. Moreover, the aging regime also emerged near the critical boundary where *α* only slightly exceeded *R*_d_, reflecting the stochastic collapse dynamics at the single-trajectory level (Fig.6(b)). Finally, in the performance-independent turnover model, the non-aging regime was observed across the entire parameter space without exception. These results indicate that the functional dependence of turnover on whole-system performance fundamentally determines whether the system maintains a finite steady-state performance or undergoes progressive collapse.

**Fig.6.**
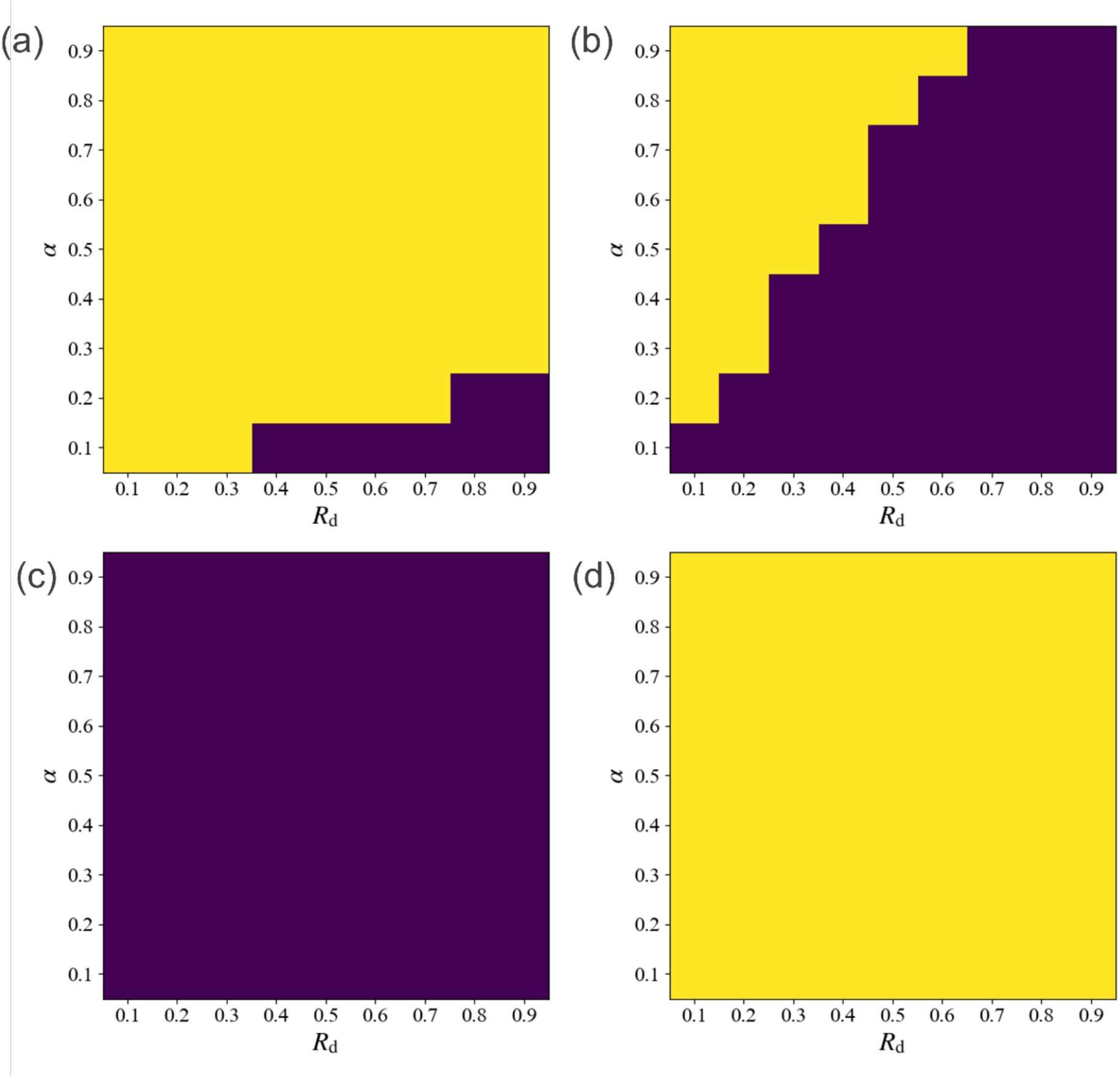
Phase diagrams showing whether a non-aging or aging regime emerges. **(a)** Model (i), **(b)** Model (ii), **(c)** Model (iii), and **(d)** the performance-independent model. The horizontal axis shows the damage rate *R*_d_, and the vertical axis shows the maximum turnover rate *α*. Both parameters are scanned from 0.1 to 0.9 in steps of 0.1. Yellow regions indicate the non-aging regime, and purple regions indicate the aging regime. Parameters are fixed at *s* = 50 for model (i) and *k* = 4 for model (iii).

## Discussion

In this study, we investigated how performance-dependent turnover influences the emergence of aging using a minimal hierarchical model linking component-level damage and turnover with whole-system performance. Across the turnover models examined here, the system behavior was consistently classified into two regimes: a non-aging regime, in which whole-system performance remained finite, and an aging regime, in which performance progressively declined toward zero. Importantly, aging emerged only when turnover depended positively on whole-system performance, indicating that the weakening of maintenance capacity at low performance constitutes a key determinant of long-term decline. Under parameter conditions belonging to the aging regime, performance loss became effectively irreversible over long timescales because the reduction of turnover increasingly limited the possibility of recovery. At the same time, individual stochastic trajectories frequently exhibited transient recovery events, showing that short-term reversibility can coexist with long-term irreversible decline.

A central mechanistic insight of the present study is that performance-dependent turnover generates a feedback structure that promotes aging. As whole-system performance decreases, the turnover rate is simultaneously reduced, thereby lowering the ability of the system to restore damaged components. This reduction further accelerates performance loss, creating a self-reinforcing process in which declining maintenance capacity amplifies the initial deterioration. Analytical calculations showed that this structural dependence alone is sufficient to generate aging dynamics in specific parameter regions, even without introducing additional biological assumptions. Moreover, numerical simulations demonstrated that stochastic fluctuations can broaden the apparent aging regime by inducing collapse events that are absent in deterministic trajectories. These findings suggest that, in biological systems where fluctuations are unavoidable, aging-like dynamics may emerge under a wider range of conditions than predicted solely from equilibrium analysis.

The emergence of aging in the present framework should not be regarded as an arbitrary consequence of parameter choice. Rather, it reflects a physically plausible asymmetry between damage generation and turnover-mediated restoration. Damage can arise rapidly, passively, and at multiple spatial locations through intrinsic physicochemical processes or environmental perturbations. In contrast, turnover is an active maintenance process that requires time, metabolic energy, molecular machinery, and spatially localized execution. Thus, even when turnover is efficient, it cannot instantaneously and globally compensate for damage occurring across spatially distributed components. This temporal, spatial, and energetic asymmetry provides a physical basis for why biological systems may be biased toward parameter regimes favoring aging. In the linear model, this tendency is analytically represented by the condition *α* < *R*_d_, whereas analogous aging regimes also emerge in the other turnover formulations. In this sense, the aging regime identified here is not merely a parameter-dependent outcome of the model, but a physically favored consequence of the constrained nature of biological maintenance.

Although aging emerged as an effectively irreversible process over long timescales, the model also revealed that performance does not necessarily decline monotonically at the level of individual stochastic trajectories. Short-term increases in performance arose when turnover transiently exceeded damage, even under parameter conditions that ultimately led to collapse. Such local recovery does not prevent long-term decline, but it indicates that aging can proceed through intermittent episodes of partial restoration superimposed on an overall downward trend. These stochastic fluctuations also produced substantial variation in the timing of collapse, leading to lifespan variability across trajectories despite identical parameter values. Thus, the model reconciles two features often observed in biological aging, namely the long-term directionality of functional decline and the presence of transient functional improvement or rejuvenation-like events.

A recent theoretical framework modeled aging as a stochastic process based on the production and removal dynamics of senescent cells and showed that their age-dependent accumulation can explain Gompertz-like increases in mortality (Karin et al., 2019). In that framework, the best-supported model was a saturating removal model, in which senescent-cell production increases with age and accumulated senescent cells slow their own removal, producing critical slowing down, persistent fluctuations, and first-passage-like mortality. This work is closely related to the present study because both emphasize turnover, stochastic fluctuations, and collapse-like transitions. However, whereas the previous framework focused on a specific aging-associated factor and mortality statistics, the present study identifies a more general dynamical structure in which component-level damage reduces whole-system performance, and reduced performance in turn weakens turnover capacity. Thus, our model directly isolates the damage–turnover balance as a minimal principle underlying aging-like decline and long-term irreversibility.

The same damage–turnover coupling can be interpreted across multiple biological scales. At the molecular level, DNA damage accumulates when damage generation exceeds repair capacity. Mitochondrial aging can also be viewed through this framework, because mitochondrial DNA mutations and structural deterioration accumulate when quality-control processes such as mitophagy fail to compensate. This imbalance may be further amplified when reduced mitochondrial performance limits the energy available for turnover. At the cellular and tissue levels, decline in stem-cell function represents a failure of turnover capacity itself, because impaired stem cells or their niches reduce the replacement of dysfunctional cells. These examples suggest that diverse aging-associated phenomena may be understood as scale-specific manifestations of the same general principle, in which functional decline emerges when damage accumulation and performance-dependent turnover become dynamically coupled.

Although the present framework provides a general description of aging arising from component-level damage and turnover, it does not directly apply to all biological systems. Importantly, this limitation does not necessarily represent a contradiction to the model, because several organisms exhibiting negligible senescence or apparent immortality rely on maintenance mechanisms operating at hierarchical levels different from those assumed here. The present model considers damage and turnover occurring among constituent components within an individual, whereas some biological systems employ turnover at the organismal or lineage level. For example, *Turritopsis dohrnii* can revert from the mature medusa stage to the polyp stage through transdifferentiation, effectively resetting accumulated damage at the organismal level (Matsumoto et al., 2019). Similarly, lineage immortality in unicellular organisms is maintained because cell division itself functions as organism-level renewal (Florea, 2017). Continuous tissue replacement in organisms such as *Hydra (Reddy et al*., *2019)* may also operate through maintenance mechanisms extending beyond simple component-level repair. These systems therefore should not be regarded as exceptions to the present framework, but rather as examples of distinct hierarchical organizations of turnover.

Another limitation of the model is that the decline in performance primarily follows exponential-like trajectories, whereas aging in many organisms is often represented by logistic behavior, in which functional decline remains slow at early stages and accelerates later in life. This difference arises because the present model was designed to explain why aging emerges, rather than to reproduce all temporal details of aging trajectories. Nevertheless, several observations suggest that the framework still captures biologically relevant features. Cellular biomarkers associated with metabolic capacity, including age-dependent decreases in NAD^+^, have been reported to exhibit approximately exponential decline (Massudi et al., 2012). Moreover, the stochastic simulations showed that random damage events can induce abrupt collapses in performance, generating trajectories resembling rapid late-life deterioration. Future extensions incorporating spatially distributed repair, finite propagation of turnover, energetic constraints, and multilevel hierarchy may therefore provide a more complete description linking the present framework with organism-level aging dynamics.

